# MiBiOmics: An interactive web application for multi-omics data exploration and integration

**DOI:** 10.1101/2020.04.24.031773

**Authors:** Johanna Zoppi, Jean-François Guillaume, Michel Neunlist, Samuel Chaffron

## Abstract

**Background:** Multi-omics experimental approaches are becoming common practice in biological and medical sciences underlying the need to design new integrative techniques and applications to enable the holistic characterization of biological systems. The integrative analysis of heterogeneous datasets generally allows us to acquire additional insights and generate novel hypotheses about a given biological system. However, it can often become challenging given the large size of omics datasets and the diversity of existing techniques. Moreover, visualization tools for interpretation are usually non-accessible to biologists without programming skills.

**Results:** Here, we present MiBiOmics, a web-based and standalone application that facilitates multi-omics data visualization, exploration, integration, and analysis by providing easy access to dedicated and interactive protocols. It implements advanced ordination techniques and the inference of omics-based (multi-layer) networks to mine complex biological systems, and identify robust biomarkers linked to specific contextual parameters or biological states.

**Conclusions:** Through an intuitive and interactive interface, MiBiOmics provides easy-access to ordination techniques and to a network-based approach for integrative multi-omics analyses. MiBiOmics is currently available as a Shiny app at https://shiny-bird.univ-nantes.fr/app/Mibiomics and as a standalone application at https://gitlab.univ-nantes.fr/combi-ls2n/mibiomics.

## Background

The holistic characterization of biological systems is extending our knowledge about the functioning of organisms and natural ecosystems. Today, their multi-omics characterization is becoming standard, thus novel methodologies and easily accessible tools are required to facilitate the study of associations and interactions within and across omics layers (e.g. (meta-)genome, (meta-)transcriptome, metabolome) and scales (e.g. cells, organs, holobionts, communities). The analysis of single omics datasets has helped to identify molecular signatures associated to phenotypes of interest. However, it usually does not allow to predict mechanisms underlying phenotypic variabilities. Although multi-omics information is not sufficient to identify causes and consequences of a biological process, it can contribute to delineate key players sustaining it [1]. Indeed, exploring a biological system across several omics layers enable to capture additional sources of variability associated with a variation of interest and potentially to infer the sequence of events leading to a specific process or state [2]. Within the last decade, multi-omics integrative approaches have been applied across various fields including microbial ecology [3], genetics [4] and personalized medicine [5]. As of today, several integrative methods have been developed, but are often specific to a given experimental design, data type or a precise biological question [6]. Considering the multiplicity of existing techniques, the selection of an appropriate workflow is challenging for biologists, especially when it comes to the representation of several system-level omics layers and its interpretation. There is a clear need for accessible (web) tools to facilitate the integration, analysis and representation of multi-omics datasets through an intuitive and guided approach. MiBiOmics aims to provide established and novel techniques to reveal robust signatures in high dimensional datasets [7] through a graphical user interface allowing to perform widely applicable multi-omics analyses for the detection and description of associations across omics layers. Available as a web-based and a stand-alone application, it gives access to several R packages and tools to help users who are not familiar with programming to load and explore their data in a simple and intuitive way. MiBiOmics allows the parallel study of up to three omics datasets, as well as the in-depth exploration of each single dataset. It also provides easy access to exploratory ordination techniques and to the inference of (multi-layer) correlation networks enabling useful dimensionality reduction and association to contextual parameters. The user can then compare results from these different approaches and cross-validate multi-omics signatures to generate confident novel hypotheses.

## Implementation

MiBiOmics is implemented in R (Version 3.6.0) as a Shiny app providing an interactive interface to perform each step of a single- or multi-omics data analysis (Figure 1). MiBiOmics is also accessible as a standalone application that can be easily installed via Conda (Version 4.6.12). The application is divided into five sections as described below:

**Figure 1.**
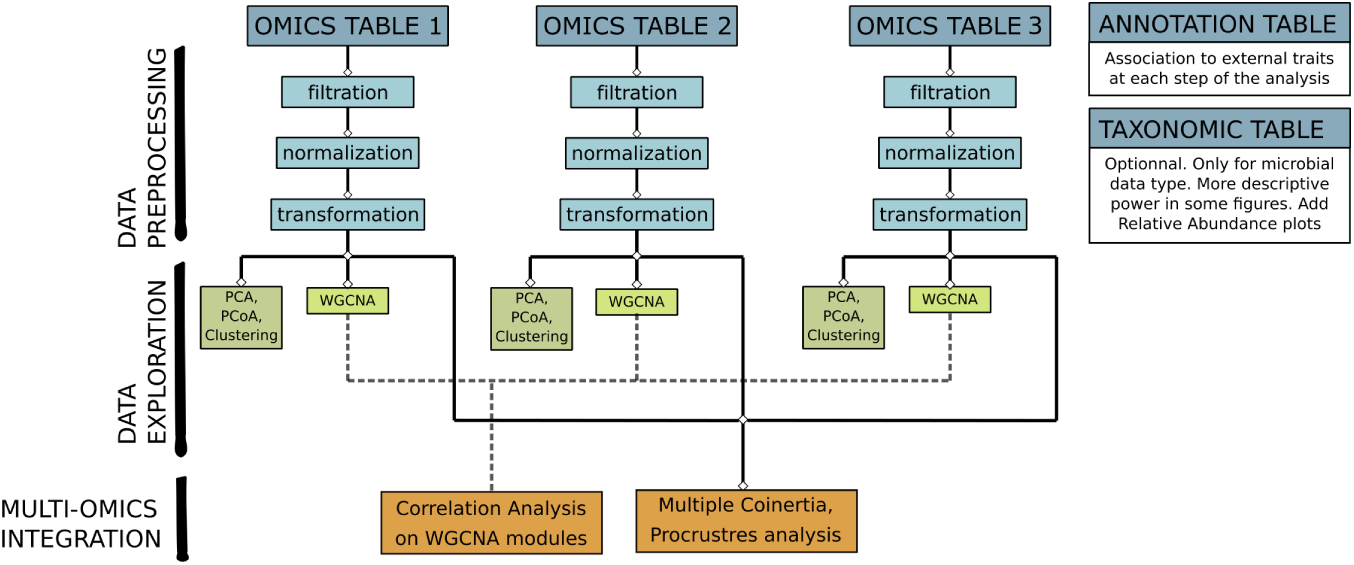
MiBiOmics framework. The MiBiOmics workflow can be divided into three main tasks: Data preprocessing, data exploration, and multi-omics integration. The data preprocessing task is dedicated to data upload, data filtration, normalization and transformation. The data exploration task implements classical clustering methods, PCA, PCoA and WGCNA correlation networks that can be applied to each omics dataset separately. Finally, the multi-omics integration task allows the user to perform multi-omics exploration, integration and analyses using ordination techniques (multiple co-inertia and Procrustes analysis), and multi-omics network inference.

### Data Upload

Within MiBiOmics, the user can upload up to three omics datasets, allowing the data exploration and network analysis of a single- or multi-omics dataset. There must be common samples between omics datasets in order to perform all analyses provided by the application. An annotation table describing external parameters (e.g. pH, site of extraction, physiological measures) needs to be provided. These parameters may be quantitative or qualitative, and available for each sample. An additional taxonomic annotations table can be uploaded when one omics table corresponds to microbial lineages (e.g. as Operational Taxonomic Units (OTUs) or Amplicon Sampling Variants (ASVs)).

Following data upload, the user can filter, normalize and transform each data matrix using common methods, such as the center log ratio (CLR) transformation to deal with the compositional nature of sequencing data, or filtration based on prevalence. In this section, it is also possible to detect and remove potential outlier samples. To allow new users to easily test the functionality of MiBiOmics, we provide two example datasets: the breast TGCA datasets from *The Cancer Genome Atlas* [8] allows to explore assocations between miRNAs, mRNAs and proteins in different breast cancer subtypes; and a dataset from the *Tara* Oceans Expeditions [9, 10] to explore prokaryotic community compositions across depth and geographic locations.

### Data Exploration

In this section, several ordination plots (Principal Component Analysis (PCA), Principal COordinates Analysis (PCoA)) [11] are dynamically produced to visualize and explore relationships between samples, and to identify main axes of variation in each dataset. When OTUs or ASVs are uploaded with their taxonomic annotations, it is possible to obtain a relative abundance plot describing the proportion of lineages at a given taxonomic level (e.g. Phylum, Family, Genus or Species) in each sample.

### Network Inference

The network inference section allows to perform a Weighted Gene Correlation Network Analysis (WGCNA[12]). Help sections are available to assist the user with parametrization, notably for optimizing the scale-free topology of the network. Here, WGCNA networks can be inferred for each uploaded omics dataset. We strongly advise users to read the WGCNA original publication and associated tutorials for this step of the analysis.

### Network Exploration

The network exploration section allows to compute and explore significant associations between subnetworks or modules (e.g. of genes, transcripts, metabolites), and communities (of lineages) delineated from each omics layer, which are denser and contain highly correlated features. Each module is associated to all external parameters provided in the annotation table and correlations are visualized as a heatmap (Figure 2A). Modules associated to parameters of interest can be further analyzed. The user can also identify which samples are contributing the most to the delineation of a specific module (Figure 2B). In case an OTUs/ASVs table is provided with taxonomic annotations, the relative abundance of lineages contributing to each module can be visualized as barplots.

**Figure 2.**
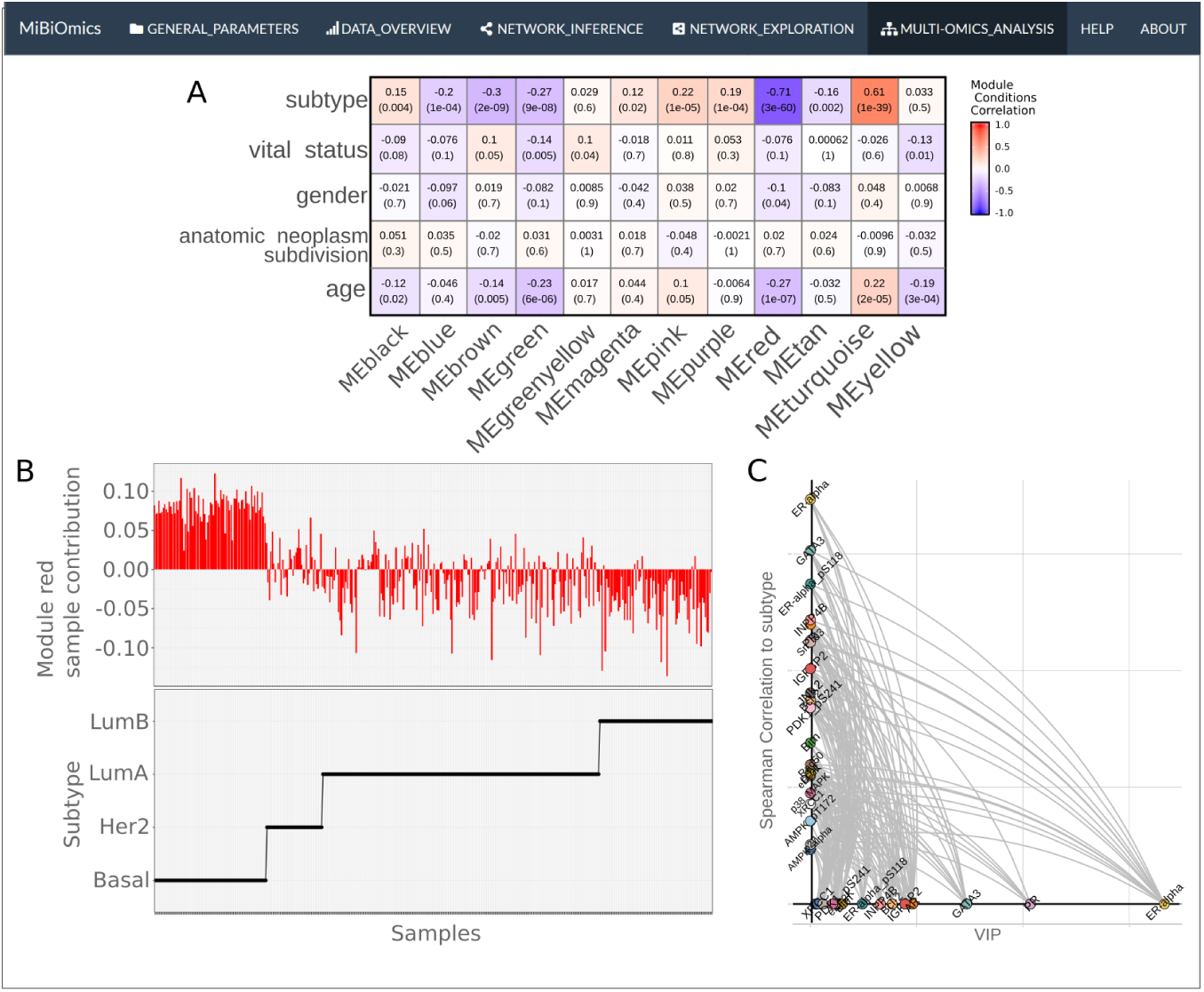
MiBiOmics networks exploration. MiBiOmics networks analysis visualizations (network exploration section) from the analysis of the The Cancer Genome Atlas Network [8] breast TGCA datasets. A. Correlation heatmap displaying associations of interest between WGCNA modules and contextual parameters. B. The upper panel indicates the contribution of each sample in the red module delineation. The lower panel indicates the corresponding subtype value for each sample. Here, Basal samples positively contributes module red, while Her2, LumA and LumB negatively contributes to module red. C. Hive plot displaying the red module’s features according to their VIP scores, correlations to the subtype parameter and their relationships.

In addition, OPLS (Orthogonal Partial Least Square) regressions [13] can be performed using a selected module components as features in order to estimate its capacity to predict a given contextual parameter, and are useful to cross-validate a module-parameter association. The results of this analysis are represented as hive plots with two axes. On the x-axis, the module features are ordered according to their Variable Importance Projection (VIP) score (a measure of their weight in the OPLS regression), while on the y-axis they are ordered according to their correlations to an external parameter of interest (Figure 2C).

### Multi-Omics Analysis

Here, MiBiOmics allows users to detect and study associations across omics datasets. Ordination methods including Procrustes analysis [11] and multiple co-inertia [14] are useful to compute and visualize the main axes of covariance, and also to extract multi-omics features driving this covariance. This central section of MiBiOmics implements an innovative approach for detecting robust links between omics layers. All modules delineated within each omics-specific network are associated to each other by directly correlating their eigenvectors. Here, the dimensionality reduction of each omics dataset through module definition ensures a small number of correlations, thereby increasing the statistical power for detecting significant associations between omics layers. For visualization, a hive plot helps summarizing significant associations between each module as a multi-layer network integrating links between omics-specific modules as well as their association to contextual parameters (traits or phenotypic characteristics). In this hive plot, each axis represents the network of a given omics layer. Corresponding modules are ordered on the axes according to their association to a contextual parameter of interest selected by the user. Modules with no significant associations are not depicted. Significant associations between omics-specific modules are represented, and individual associations between modules can also be visualized as heatmaps and dataframe. Conveniently, the user can also select modules of interest to investigate pairwise correlations between modules’ features and delineates group of modules associated together and to an external parameter.

Importantly, all figures generated by the application (PCA, PCoA, relative abundance plots, WGCNA outputs, hive plots, multiple co-inertia, Procrustes plots, correlograms, bipartite networks) can be downloaded (as svg or pdf files), and if aWpplicable, supplementary information provided as a csv file (WGCNA modules information, eigenvalues and coinertia drivers).

## Results and Discussion

MiBiOmics enables the exploration, integration, analysis and visualization of up to three omics datasets. Through the primary exploration of a dataset, the inference of biological networks and the extraction of multi-omics associated features, the application provides a ready-to-use analysis pipeline to interactively explore sources of variability and variables of interest in a given biological dataset, as well as associations between multi-omics features in holistic studies.

The inference of networks from omics features is useful to represent and model the complex architecture of putative interactions in biological systems. In addition, networks provide a way to reduce the dimensionality of a dataset by delineating cohesive groups of co-varying, often functionally related features, that can then be associated to contextual or phenotypic characteristics of interest [1]. A key functionality of MiBiOmics is the multi-omics adaptation of WGCNA [12] to explore association across omics datasets via a network-based approach. As shown in Figure 2A, the interface provides the ability to interactively probe associations across omics layers of different breast cancer subtypes [8] within each network and their association to patient parameters. The original WGCNA outputs are provided by the application to deepen the analysis between modules and external parameters (Figure 2B). In addition, we provide the user with the possibility to perform an OPLS regression for modules of interest to evaluate the robustness of these variables to predict a given trait or phenotype. Figure 2C is an example of an OPLS regression using WGCNA module variables as features. On the x-axis the features of the red module are ordered according to their VIP score (their importance for the module), and on the y-axis according to their correlation to the subtype parameter. This figure highlights how central features of a WGCNA module relate to an external parameter.

The exploratory multi-omics analysis allows to study the main axes of covariance across omics profiles and give the ability to discover and select variables implicated in an association between omics datasets. The concomitant application of (multiple) co-inertia (Figure 3A) and/or Procrustes ordination techniques with the exploration of multi-omics correlations between WGCNA modules of distinct omics layer (Figure 3B), provides a complementary vision of multi-omics relationships. The MiBiOmics interface allows to explore WGCNA modules of interest to directly infer significant associations between features from distinct omics layers (Figure 3C). In a multi-omics adaptation, WGCNA can be used to delineate a group of modules associated together and to a parameter to interest and extract features of different omics nature but related to each other. While an interactive version of WGCNA already exists [15], MiBiOmics goes beyond by providing a multi-omics strategy to identify correlated modules across omics layers and generate novel hypotheses. Associating modules across different datasets has already been performed in the original WGCNA article [12] and reproduced in several studies. For example, the overlap of modules between transcriptional profiles of different tissue [16] was assessed, as well as a comparison between proteomics and gene expression profile of modules in a cohort of alzheimer patients [17]. In both cases, the association between modules was determined by overlapping identical features (e.g. same genes in a given reference genome) within each module, a method which is not applicable when omics datasets do not contain similar data types or refer to the same biological system. In MiBiOmics, we enable the inference of relationships between omics layers within an entire biological system (e.g. holobiont) or ecosystem (e.g. the plankton), which makes it more widely applicable and especially suited for omics-based environmental studies.

**Figure 3.**
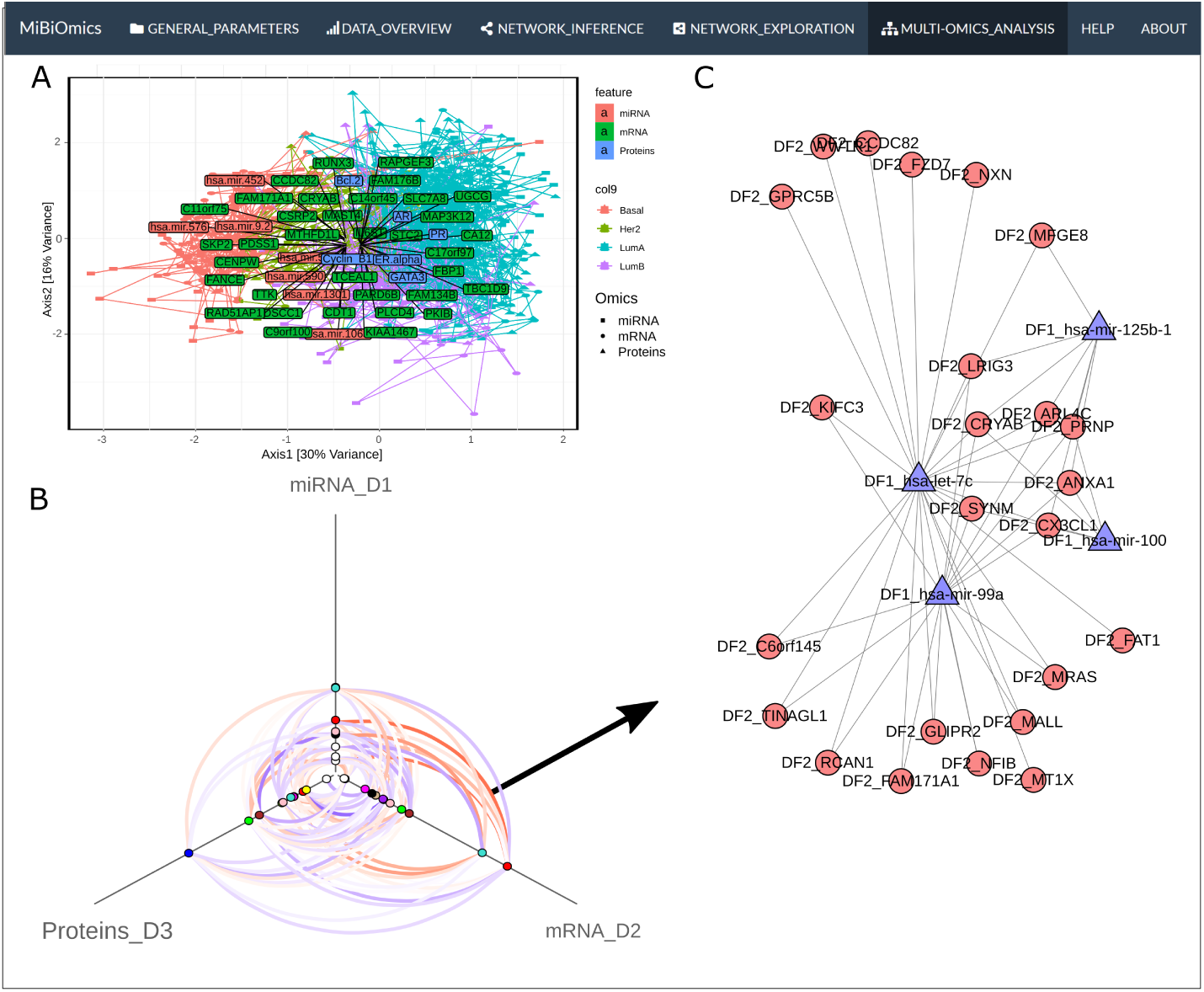
MiBiOmics multi-omics integration. MiBiOmics multi-omics integration visualizations (multi-omics integration section) from the analysis of the The Cancer Genome Atlas Network [8] breast TGCA datasets. A. A multiple co-inertia plot integrating 3 omics layers and extracted miRNA, mRNA and protein drivers. B. Hive plot displaying modules of each omics network and their associations. C. Bipartite network between mRNA features of the red mRNA module and miRNA features of the red miRNA module (Spearman Correlation > 0.35).

While there is a growing need for multi-omics analysis tools, a few applications have been developed to fulfill the need of integrating and mining these complex and heterogeneous datasets. Several methods such as MONGKIE [18], are based on prior knowledge and integrate data by projecting them on known metabolic networks and biological pathways. More generally, existing multi-omics pipelines are focusing on certain data types (Metabolomics with MetaboAnalyst [19]) or on disease-related mechanisms (MergeOmics [20]). More widely applicable methods exist, such as the R package mixOmics [21] that provides several semi-supervised methodologies often based on ordination techniques. We compared methods integrated in MiBiOmics (see Supplementary information for details) to the mixOmics DIABLO methodology [22]. Only few multi-omics features associated to breast cancer subtype in the TGCA dataset were extracted by all three methods (n=32, Figure 4A and Supplementary table 1). Both methods integrated in MiBiOmics (i.e. multiple co-inertia and multi-omics WGCNA) and DIABLO extracted mostly distinct features (Figure 4A) underlying the probable complementarity of these multi-omics integrative strategies. Scores attributed by each method to the common set of extracted features were also dissimilar (Figure 4BCD and Supplementary table 1). This may be explained by the fact that these methods implement fundamentally different approaches to features extraction and selection, which confirms the complementary nature of each analysis. For comparing the predictive power of models integrating features extracted by each method, we performed Sparse Partial Least Square Discriminant Analysis (sPLS-DA) and computed the corresponding mean AUC scores (Figure 4A and Supplementary Figure 1). All models can be considered to be highly predictive of the cancer subtype phenotypes with the miBiOmics multiomics WGCNA methodology obtaining the highest AUC score (AUC=0.9952), while the multiple coinertia analysis performed very well too (AUC=0.9903). Features extracted by the DIABLO method from mixOmics obtained the lowest score (AUC=0.9728) but remained highly predictive. Generally, these methods may benefit from an enrichment method applied to the list of extracted drivers [14]. Overall, MiBiOmics provides two complementary methods to extract associated variables between omics layers and in relationship with a trait of interest. While it is well suited to generate new hypotheses about molecular processes, it can not infer causal mechanisms between omics features and phenotypes, which would require experimental validations but that can actually be guided by MiBiOmics results.

**Figure 4.**
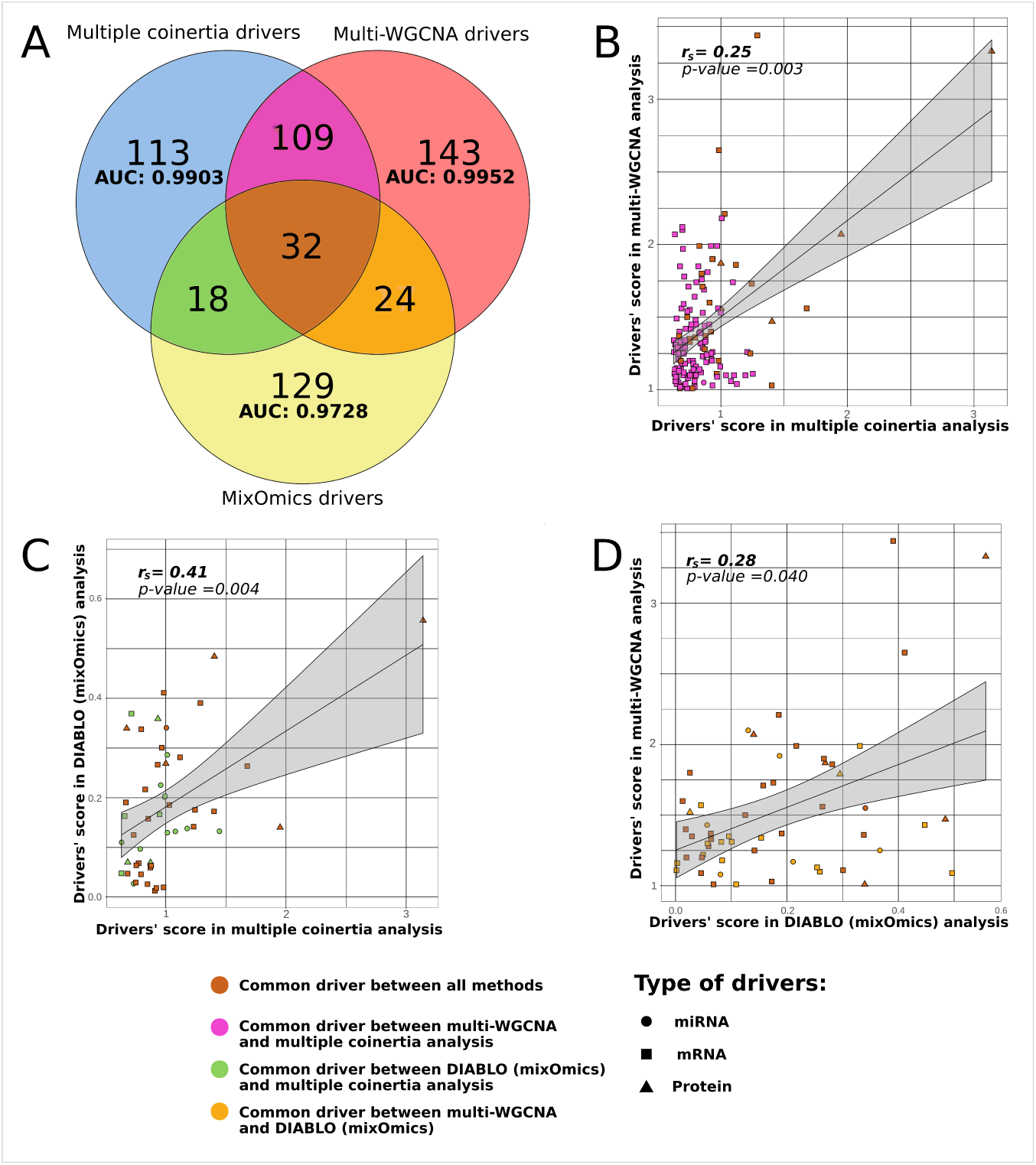
MiBiOmics and mixOmics comparison. A. Venn diagram displaying common and distinct features extracted by DIABLO (mixOmics), multiple coinertia and multi-omics WGCNA (MiBiOmics). Area Under the Curve (AUC) scores were computed to compare features selection and model performance of each method. B. Selected features’ weights comparison between multiple coinertia and multi-WGCNA on their common subset of features. C. Selected features’ weights comparison between multiple coinertia and DIABLO (mixOmics) on their common subset of features. D. Selected features’ weights comparison between DIABLO (mixOmics) and multi-WGCNA method on their common subset of features.

In addition to provide an exploratory and integrative framework for multi-omics studies, MiBiOmics distinguishes itself by providing a powerful dimensionality reduction and unsupervized method combining both ordination and graph-based techniques, which enables to study complex biological systems as a whole. Importantly, it also integrates contextual information by linking multi-omics signatures to qualitative and quantitave contextual parameters.

## Conclusion

MiBiOmics is an interactive web-based (and standalone) application to easily and dynamically explore associations across omics datasets. Through an innovative network-based integrating strategy, it can help biologists to identify potential mechanisms of interactions and generate novel hypotheses. The core of the application lies behind the reduction of dimensionality across omics datasets to efficiently link them at the molecular level, and to identify biomarkers associated with a given trait or phenotype. The MiBiOmics pipeline facilitates the exploration, integration and analysis of multi-omics datasets to a broad audience by providing scientists a powerful way to predict and explore putative molecular mechanisms underlying complex phenotypes across a wide range of biological scales and systems.

## Availability and requirements

**Project name:** MiBiOmics

**Project home page:** https://gitlab.univ-nantes.fr/combi-ls2n/mibiomics

**Operating system(s):** Platform independent

**Programming language:** R

**Other requirements:** for the local installation Conda 4.6.12 or Docker.

**License:** AGPL-3

**Any restrictions to use by non-academics:** No restrictions.

## List of abbreviation

ASV: Amplicon Sequence Variant.
AUC: Area Under the Curve.
DIABLO: Data Integration Analysis for Biomarker discovery using Latent variable approaches for ‘Omics studies.
PCA: Principal Component Analysis.
PCoA: Principal COordinates Analysis.
OPLS: Orthogonal Partial Least Square.
OPLS-DA: Orthogonal Partial Least Square Discriminant Analysis.
OTU: Operational Taxonomic Unit.
VIP: Variable Importance Projection.
WGCNA: Weighted Gene Correlation Network Analysis.

## Ethics approval and consent to participate

Not applicable.

## Consent for publication

Not applicable.

## Availability of data and materials

The datasets provided as example within MiBiOmics application are available in the data repository, at https://gitlab.univ-nantes.fr/combi-ls2n/mibiomics.

## Authors’ contribution

JZ and SC participated equally in the design and development of MiBiOmics as a standalone web-application, and writing the manuscript. MN contributed to the writing of the manuscript. JFG was responsible for the distribution and maintenance of MiBiOmics web-server at https://shiny-bird.univ-nantes.fr/app/Mibiomics. All authors read and approved the final version of the manuscript.

## Acknowledgements

The authors wish to thank Mélanie Fouesnard, Damien Eveillard, Philippe Bordron, Gaëlle Boudry, Catherine Michel and Simon Beck for their useful feedback while testing and using the application. We also thank the bioinformatics core facility of Nantes (BiRD - Biogenouest) for providing computing resources and support.

## References

1. Hasin, Y., Seldin, M., Lusis, A.: Multi-omics approaches to disease. Genome Biology 18(1), 1–15 (2017). doi:10.1186/s13059-017-1215-1. Hasin, Yehudit, 2017, Multi-omics

2. Li, Y., Wu, F.-X., Ngom, A.: A review on machine learning principles for multi-view biological data integration. Briefings in Bioinformatics 19 (October 2016), 113 (2016). doi:10.1093/bib/bbw113

3. Heintz-Buschart, A., May, P., Laczny, C.C., Lebrun, L.A., Bellora, C., Krishna, A., Wampach, L., Schneider, J.G., Hogan, A., De Beaufort, C., Wilmes, P.: Integrated multi-omics of the human gut microbiome in a case study of familial type 1 diabetes. Nature Microbiology 2(1), 1–12 (2016). doi:10.1038/nmicrobiol.2016.180

4. Zhang, B., Gaiteri, C., Bodea, L.G., Wang, Z., McElwee, J., Podtelezhnikov, A.A., Zhang, C., Xie, T., Tran, L., Dobrin, R., Fluder, E., Clurman, B., Melquist, S., Narayanan, M., Suver, C., Shah, H., Mahajan, M., Gillis, T., Mysore, J., MacDonald, M.E., Lamb, J.R., Bennett, D.A., Molony, C., Stone, D.J., Gudnason, V., Myers, A.J., Schadt, E.E., Neumann, H., Zhu, J., Emilsson, V.: Integrated systems approach identifies genetic nodes and networks in late-onset Alzheimer’s disease. Cell 153(3), 707–720 (2013). doi:10.1016/j.cell.2013.03.030. NIHMS150003

5. Chen, R., Mias, G.I., Li-Pook-Than, J., Jiang, L., Lam, H.Y.K., Chen, R., Miriami, E., Karczewski, K.J., Hariharan, M., Dewey, F.E., Cheng, Y., Clark, M.J., Im, H., Habegger, L., Balasubramanian, S., O’Huallachain, M., Dudley, J.T., Hillenmeyer, S., Haraksingh, R., Sharon, D., Euskirchen, G., Lacroute, P., Bettinger, K., Boyle, A.P., Kasowski, M., Grubert, F., Seki, S., Garcia, M., Whirl-Carrillo, M., Gallardo, M., Blasco, M.A., Greenberg, P.L., Snyder, P., Klein, T.E., Altman, R.B., Butte, A.J., Ashley, E.A., Gerstein, M., Nadeau, K.C., Tang, H., Snyder, M.: Personal omics profiling reveals dynamic molecular and medical phenotypes. Cell 148(6), 1293–1307 (2012). doi:10.1016/j.cell.2012.02.009. 1011.1669v3

6. Paliy, O., Shankar, V.: Application of multivariate statistical techniques in microbial ecology. Molecular Ecology 25(5), 1032–1057 (2016). doi:10.1111/mec.13536. 15334406

7. Guidi, L., Chaffron, S., Bittner, L., Eveillard, D., Marin, M., Roscoff, S.B.D.: Plankton networks driving carbon export in the oligotrophic ocean 532(7600), 465–470 (2016). doi:10.1038/nature16942.Plankton

8. Cancer Genome Atlas Network: Comprehensive molecular portraits of human breast tumours. Nature 490(7418), 61–70 (2012). doi:10.1038/nature11412

9. Sunagawa, S., Coelho, L.P., Chaffron, S., Kultima, J.R., Labadie, K., Salazar, G., Djahanschiri, B., Zeller, G., Mende, D.R., Alberti, A., Cornejo-Castillo, F.M., Costea, P.I., Cruaud, C., D’Ovidio, F., Engelen, S., Ferrera, I., Gasol, J.M., Guidi, L., Hildebrand, F., Kokoszka, F., Lepoivre, C., Lima-Mendez, G., Poulain, J., Poulos, B.T., Royo-Llonch, M., Sarmento, H., Vieira-Silva, S., Dimier, C., Picheral, M., Searson, S., Kandels-Lewis, S., Bowler, C., de Vargas, C., Gorsky, G., Grimsley, N., Hingamp, P., Iudicone, D., Jaillon, O., Not, F., Ogata, H., Pesant, S., Speich, S., Stemmann, L., Sullivan, M.B., Weissenbach, J., Wincker, P., Karsenti, E., Raes, J., Acinas, S.G., Bork, P.: Ocean plankton. Structure and function of the global ocean microbiome. Science (New York, N.Y.) 348(6237), 1261359 (2015). doi:10.1126/science.1261359. NIHMS150003

10. Mariette, J., Villa-Vialaneix, N.: Unsupervised multiple kernel learning for heterogeneous data integration. Bioinformatics 34(6), 1009–1015 (2018). doi:10.1093/bioinformatics/btx682

11. Dixon, P.: VEGAN, a package of R functions for community ecology. Journal of Vegetation Science 14(6), 927–930 (2009). doi:10.1111/j.1654-1103.2003.tb02228.x

12. Langfelder, P., Horvath, S.: WGCNA: An R package for weighted correlation network analysis. BMC Bioinformatics (2008). doi:10.1186/1471-2105-9-559

13. Wehrens, R., Bjørn-Helge, M.: The pls Package: Principal Component and Partial Least Squares Regression in R. Journal of Statistical Software 18(2) (2007). doi:10.18637/jss.v018.i02

14. Meng, C., Kuster, B., Culhane, A.C., Gholami, A.M.: A multivariate approach to the integration of multi-omics datasets. BMC Bioinformatics 15(1), 1–13 (2014). doi:10.1186/1471-2105-15-162

15. Sundararajan, Z., Knoll, R., Hombach, P., Becker, M., Schultze, J.L., Ulas, T.: Shiny-Seq: advanced guided transcriptome analysis. BMC research notes 12(1), 432 (2019). doi:10.1186/s13104-019-4471-1

16. Xiao, X., Moreno-Moral, A., Rotival, M., Bottolo, L., Petretto, E.: Multi-tissue Analysis of Co-expression Networks by Higher-Order Generalized Singular Value Decomposition Identifies Functionally Coherent Transcriptional Modules. PLoS Genetics 10(1) (2014). doi:10.1371/journal.pgen.1004006

17. Seyfried, N.T., Dammer, E.B., Swarup, V., Nandakumar, D., Duong, D.M., Yin, L., Deng, Q., Nguyen, T., Hales, C.M., Wingo, T., Glass, J., Gearing, M., Thambisetty, M., Troncoso, J.C., Geschwind, D.H., Lah, J.J., Levey, A.I.: A Multi-network Approach Identifies Protein-Specific Co-expression in Asymptomatic and Symptomatic Alzheimer’s Disease. Cell Systems 4(1), 60–724 (2017). doi:10.1016/j.cels.2016.11.006

18. Jang, Y., Yu, N., Seo, J., Kim, S., Lee, S.: MONGKIE: An integrated tool for network analysis and visualization for multi-omics data. Biology Direct 11(1), 1–9 (2016). doi:10.1186/s13062-016-0112-y

19. Chong, J., Soufan, O., Li, C., Caraus, I., Li, S., Bourque, G., Wishart, D.S., Xia, J.: MetaboAnalyst 4.0: Towards more transparent and integrative metabolomics analysis. Nucleic Acids Research 46 (W1), 486–494 (2018). doi:10.1093/nar/gky310

20. Shu, L., Zhao, Y., Kurt, Z., Byars, S.G., Tukiainen, T., Kettunen, J., Orozco, L.D., Pellegrini, M., Lusis, A.J., Ripatti, S., Zhang, B., Inouye, M., Mäkinen, V.P., Yang, X.: Mergeomics: Multidimensional data integration to identify pathogenic perturbations to biological systems. BMC Genomics 17(1), 1–16 (2016). doi:10.1186/s12864-016-3198-9

21. Rohart, F., Gautier, B., Singh, A., Lê Cao, K.A.: mixOmics: An R package for ‘omics feature selection and multiple data integration. PLoS Computational Biology 13(11), 1–19 (2017). doi:10.1371/journal.pcbi.1005752

22. Singh, A., Gautier, B., Shannon, C.P., Vacher, M., Rohart, F., Tebbutt, S.J., Lê Cao, K.A.: DIABLO: an integrative approach for identifying key molecular drivers from multi-omics assays. Bioinformatics 35(17), 3055–3062 (2019)

