## Supplementary information for "MiBiOmics: An interactive web application for multi-omics data exploration and integration"

Details about multi-omics data analysis methodologies we compared in the article (Figure 4) are described below. Both methods implemented in MiBiOmics (multi-WGCNA and multiple co-inertia) were compared to DIABLO [1] integrated in the mixOmics R package.

**mixOmics DIABLO**

We performed a DIABLO analysis on the whole breast TCGA dataset to extract multi-omics features associated to the tumor subtype parameter. Following the DIABLO tutorial [1], we choose a design where all the omics blocks (mRNA, miRNA and proteins) are connected with a link of 0.1. For selecting the final model we choose the centroid distance with 4 components, and identified an optimum number of extracted features per components using the function *tune.block.plsda*.

A total of 203 non-redundant features were selected using this protocol, which we compared with both methods implemented in MiBiOmics.

**MiBiOmics multi-WGCNA**

For the integration of multi-omics datasets, MiBiOmics allows the inference of multi-layer networks based on the WGCNA methodology developed by Langfelder and Horvath in 2008 [2]. This multi-layer network is built by detecting significant associations between WGCNA subnetworks or modules delineated for each omics dataset. In addition, association to contextual information is also integrated by detecting modules of the multi-layer network significantly associated to a given trait or phenotype. To extract multi-omics features associated to a contextual parameter of interest (here the tumor subtype in the TGCA dataset), the following protocol was implemented:

- WGCNA signed networks are inferred for all omics datasets (miRNA, mRNA and protein datasets). Here, we used a biweight midcorrelation (or bicor), and choose soft powers of 16, 8 and 10 with a minimum module size of 4, 6 and 4 for the miRNA, mRNA and protein datasets, respectively. For these parametrization steps (soft power and minimum module size), we strongly advise users to follow protocols and instructions associated to the WGCNA article [2].
- Modules associated to our trait of interest (tumor subtype) were selected based on the Spearman correlation (and associated p-value) between the parameter and the modules eigenvalue (abs(cor.) > 0.5 and p-value < 0.001). Based on these criteria, three modules were selected: the mRNA red and turquoise modules, and the protein green module.
- Starting from this first set of modules we delineated a group of modules significantly associated together. The hive plot in the MiBiOmics 'multi-omics analysis' section allows to visualize how eigenvalues of each module correlate to each other across omics layers. This step allows to detect significant associations between modules and thus between omics layers. Here, we selected modules associated to the first set of modules directly associated to the trait of interest (the red and turquoise from the mRNA network, and the green from the protein dataset; Spearman abs(cor.) > 0.5 and p-value < 0.001. Using this procedure, we obtained a multi-layer network or network of modules associated to a given trait. At this stage, the following additional modules were selected: blue and turquoise from the miRNA network, brown, red and turquoise from the mRNA network and blue and green from the protein network.
- For each module, the list of features, their VIP scores, correlations to the subtype parameter and associated p-value were downloaded via the '*Network Exploration*' tab, after setting the appropriate number of components for each O-PLS (the optimum number of components is

identified by the first minimum local on the Root Mean Square Error of Prediction (RMSEP) plot.
- Given some modules may contain many features, we selected these features weighted by their importance in the module (based on the VIP score), and their association to the parameter of interest (tumor subtype). Here, we selected only features that obtained a VIP score above 1 and an associated p-value below 0.05.

Using this protocol, 308 features were selected across the mRNA, miRNA and protein datasets to be significantly related together and/or associated to the tumor subtype.

**MiBiOmics multiple co-inertia**

Using the TGCA multi-omics dataset, we performed a multiple co-inertia as implemented in MiBiOmics with the ade4 R package, and extracted drivers on the first axis of co-variance (we selected the first axis of the multiple co-inertia along which samples were ordered according to their respective subtype). These drivers or features are ranked according to how much they participate to the co-variance on this axis. Here, we selected the top 30% features with the highest absolute score in the first axis of the total covariance.

Following this procedure, a total of 272 multi-omics features were extracted.

**Comparing the predictive power of each method**

In order to compare the capacity of these methods to extract features associated to a parameter of interest we performed a Sparse Partial Least Squares Discriminant Analysis (sPLS-DA) using each method features associated to the tumor subtypes using the mixOmics *plsda* function. The appropriate number of components was chosen using the recommended value of the *perf* mixOmics function and the more accurate distance metric (the selected number of components for the sPLS-DA was 6, 7 and 3 for the multiple co-inertia, multi-WGCNA and DIABLO features, respectively). The AUC was computed and ROC curves were plotted for each sPLS-DA (Figure S1) to estimate and compare the predictive power of each method according to the tumor subtype parameter. The AUC indicated a strong predictive power for all three methodologies (DIABLO-AUC = 0.973, multi-WGCNA-AUC = 0.999, multiple co-inertia-AUC = 0.990) but using distinct extracted features.

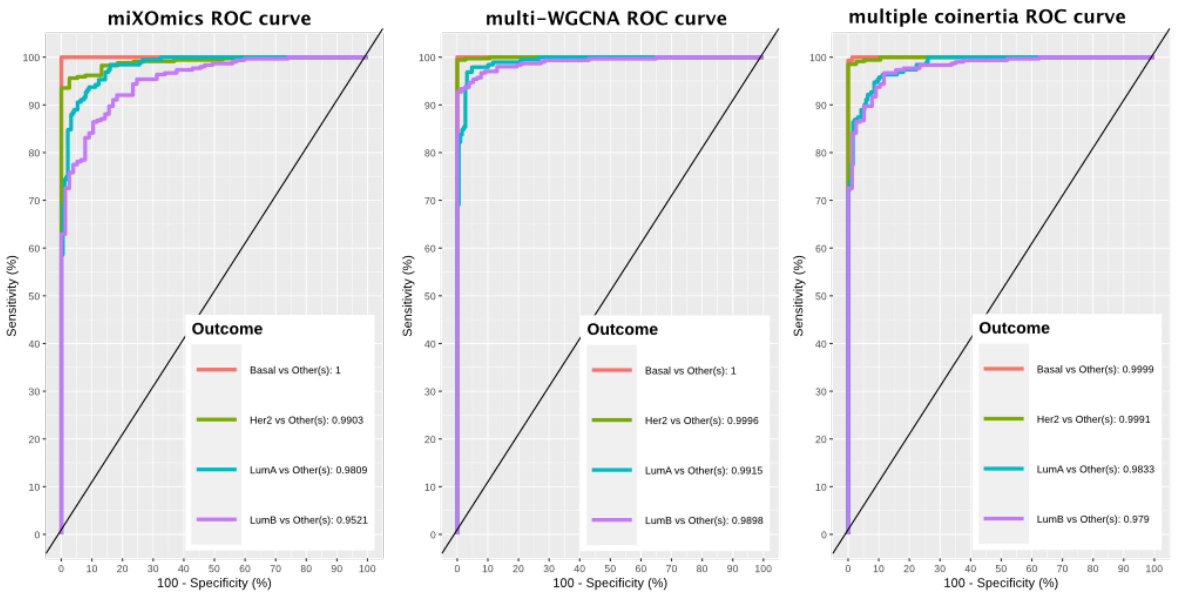


**Figure S1 : Predictive power of DIABLO (mixOmics), multi-WGCNA (MiBiOmics) and multiple coinertia (MiBiOmics) for the tumor subtype parameter of the TCGA dataset.** ROC curves and AUC were obtained using sPLS-DA models built using extracted features associated to the tumor subtype parameter by the three methods (DIABLO from mixOmics, multi-WGCNA and multiple co-inertia from miBiOmics).

### References


1. Singh, A., Gautier, B., Shannon, C.P., Vacher, M., Rohart, F., Tebbutt, S.J., Lê Cao, K.A.: DIABLO: anintegrative approach for identifying key molecular drivers from multi-omics assays. Bioinformatics35(17),3055–3062 (2019)

2. Langfelder, P., Horvath, S.: WGCNA: An R package for weighted correlation network analysis. BMCBioinformatics (2008). doi:10.1186/1471-2105-9-559
